# Principles and best practices of optimizing a micromirror-based multicolor TIRF microscopy system

**DOI:** 10.1101/2023.03.27.534330

**Authors:** Kaley McCluskey, Nynke H. Dekker

**Affiliations:** Department of Bionanoscience, Delft University of Technology. Building 58, van der Maasweg 9, 2629HZ Delft, Netherlands

**Author notes:** Corresponding author. E.

**Keywords:** single-molecule fluorescence microscopy, (co)localization microscopy, TIRF, micromirror, CoSMoS

## Abstract

TIRF (Total Internal Reflection Fluorescence) microscopy is a powerful tool for measuring the intra- and intermolecular dynamics of fluorescently-labeled single molecules. As TIRF measurements move to more complex biological systems with more fluorescent probes, the multi-band-pass dichroic that separates excitation from emission becomes limiting for the microscope’s detection efficiency. To avoid this, multicolor colocalization-based experiments can employ “micromirror” (mm)TIRF, which replaces the dichroic with two 45°-angled rod mirrors that control the TIR excitation beam(s). Whereas a dichroic spectrally separates excitation and emission wavelengths, the micromirrors act to spatially separate the excitation beams from the collected emission photons within the objective lens itself. Comprehensive control of the TIR beam in mmTIRF can yield excellent signal to noise, and hence data quality, but at the price of increased optical complexity. Here, we introduce the theory behind these additional optical components and provide practical advice from our experience on the best way to set up, align, optimize, and maintain a mmTIRF instrument. We also demonstrate the practical effects of small misalignments to illustrate both the optimized signal quality and the degree of accuracy required to achieve it. We hope that this guide increases the accessibility of this type of instrumentation and helps researchers use it to produce data of the highest quality possible.

## 1. Introduction

Total Internal Reflection Fluorescence (TIRF) microscopy is one of the most powerful tools in the single-molecule biophysical “toolkit” [1]. TIRF is widely used to study unsynchronized intra- and intermolecular interactions between fluorescently-labeled biological molecules in real time, including RNA folding, transcription, DNA replication, macromolecular complex assembly, and more.

TIRF uses the total internal reflection of a laser outside a sample chamber to illuminate a thin region near the surface with the evanescent field. This effectively excludes background fluorescence from the bulk of the sample and thereby contributes to a sufficiently high signal to noise ratio (SNR) to detect single molecules over a wide field of view. The TIRF excitation laser can be directed through a prism placed on top of the sample chamber or through the objective [1]. “Objective-type” TIRF can achieve better SNR than “prism-type” TIRF, both because no signal is lost passing through the medium to the objective and because the effective numerical aperture (NA) is limited only by the NA of the objective (1.49-1.7 for high-end TIRF objectives) and not by the refractive index of the sample, as in prism-type TIRF. (It is worth noting that going beyond NA = 1.5 for TIRF is not necessarily desirable. The non-glass cover slides and high-refractive-index immersion oil required for index matching present practical difficulties, and at the incident angles possible with NA greater than 1.7, the shallow penetration depth of the evanescent field, about 50 nm, may become limiting.) Objective-type TIRF also leaves the space above the sample available for combination with other techniques like magnetic force spectroscopy [2]. Multicolor detection can be performed with a single excitation laser when experiments involve FRET [2, 3]. However, colocalization microscopy—measurements that use the colocalization between two dyes to report on the formation or dynamics of a macromolecular complex—requires direct excitation of multiple dyes at distinct wavelengths. Objective-type TIRF typically requires a dichroic mirror to spectrally separate these excitation laser lines from the emitted fluorescence.

There is an extensive literature of high-resolution, two-color single-molecule colocalization and tracking measurements using dichroic-based objective-type TIRF [4-7]. However, as the number of fluorescence probes increases, the multi-band-pass dichroics required to separate three or more laser lines from the fluorescence emission greatly reduce the spectrum available to transmit the emitted photons. To compensate for this, an alternative to the dichroic called micromirror (mm)TIRF was developed in 2006 [8]. mmTIRF is closely associated with “Colocalization Single-Molecule Spectroscopy” (CoSMoS) as a technique[9], so we will discuss both in turn.

Micromirror TIRF [8] replaces the dichroic with a pair of micromirrors (2-3 mm diameter rod mirrors with 45º angled faces) to separate excitation from fluorescence (Figure 1a). mmTIRF improves the SNR of the system in two ways. First, by avoiding the dichroic, there is no loss of fluorescence signal around the excitation laser wavelengths, which is especially relevant for applications where many wavelengths are being used simultaneously. Second, the “output” micromirror directs the totally internally reflected output beam away from the detection optics and avoids scatter into the excitation pathway, lowering the noise.

**Figure 1.**
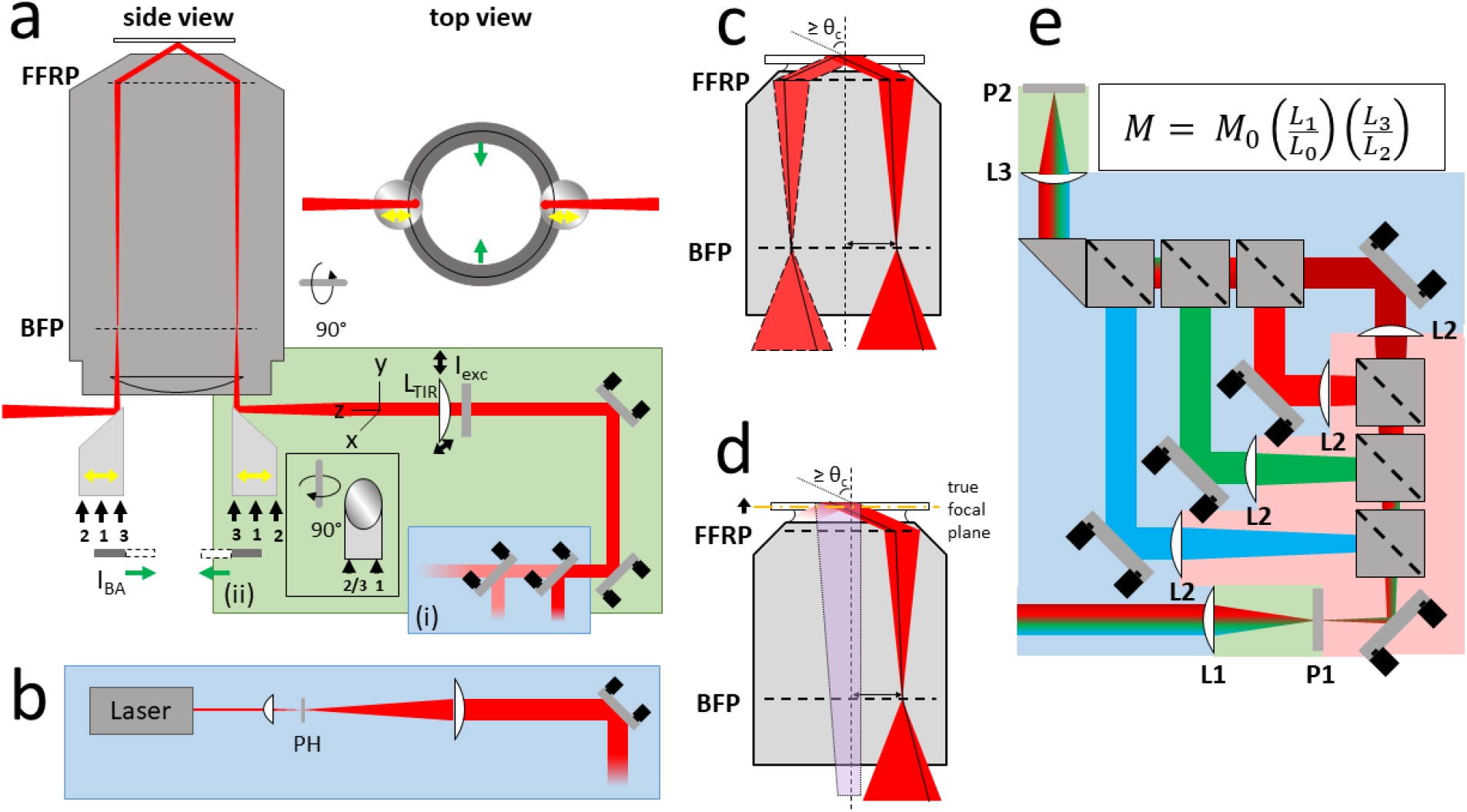
Overview of micromirror TIRF setup. (a) Schematic of the mechanical degrees of freedom in the excitation pathway. Box (i) (blue): uncoupled degrees of freedom that affect only one excitation laser. Each laser is coaligned with the others using a long-pass combining dichroic mirror on a three-point kinematic mount. The positioning of the combining dichroic affects only the reflected laser. Box (ii) (green): coupled degrees of freedom that affect all excitation lasers symmetrically. Two periscope mirrors (far right) elevate the lasers from the level of the table to that of the micromirrors, and are used to align them to the optic axis. An iris that controls the size of the TIRF beam, I_exc_, precedes the TIRF lens (L_TIR_), which is mounted in an XY kinematic holder. The micromirrors (center, below the objective barrel) are controlled by three screws that adjust their tip and tilt (numbered 1-3; also shown in the inset with a 90º rotation). The mirrors translate along the optic axis (yellow arrows). The objective is shown as a rough schematic with the effective (not actual) back focal plane (BFP) and front focal reference plane (FFRP) indicated. An adjustable iris (I_BA_: grey and dashed rectangles, green arrows) restricts the aperture. The micromirrors and iris are also show in top view (right) to illustrate that the iris just cuts the edges of the micromirrors out of the aperture. (b) Schematic beam expander for mmTIRF. A laser beam is incident on a short-focal-length, aspheric lens. A pinhole (d = 20-50 μm) is optionally placed in the Fourier plane to perform low-pass filtering. The beam is then expanded to approximately 15 mm diameter and collimated with a second lens one focal length from the pinhole. The collimated beam is coaligned with the other excitation lasers using a long-pass dichroic mirror on a three-point kinematic mount. (c) Simplified representation of the mapping of an excitation beam focused in the back focal plane (BFP) and displaced from center to a collimated TIRF footprint at angle θ_c_ at the sample interface. The solid internal lines illustrate the path of a narrower cone of light incident at an angle, resulting in a net displacement of the footprint. (d) Schematic illustrating the effect of defocus on both the positioning of the TIRF footprint and the resultant image. The correct focal plane is illustrated schematically in yellow; the sample is shown focused above the design working distance. The TIRF footprint is displaced from the optic axis, and the detected fluorescence (purple shape with dashed outline) is converging instead of collimated, so the primary image will not form one focal length from the tube lens. (e) Imaging pathway designed to visualize four color channels on quadrants of one camera. The color code indicates regions where the wave packet is collimated (blue), converging (green), or diverging (pink). Collimated light emerging from the back aperture of the objective is incident on the tube lens L_1_, which forms an image in the primary imaging plane, P_1_. An aperture is placed in P_1_ to define an image one quarter the width of the camera chip. The signal propagates past P_1_ and encounters three long-pass dichroics (grey boxes), which split the signal into four color channels. A separate lens L_2_ is placed one focal length away from P_1_ in each channel. Mirrors in each channel are used to offset the four images from one another before they are recombined using three more dichroics, then focused by lens L_3_, placed one focal length from the camera sensor in the secondary imaging plane, P_2_. The total magnification of this system depends on the magnification M_0_ of the objective, the ratio of L_1_ to the design tube length of the objective, L_0_, and the ratio of L_3_ to L_2_.

The CoSMoS experimental format monitors colocalization of multiple fluorescently-labeled “probe” molecules, usually proteins, with “target” molecules, usually DNA or RNA, immobilized on a surface. CoSMoS has been used to monitor the assembly and activation of the spliceosome [9-11], reconstitute the licensing and assembly of the CMG replicative helicase [12-14], and characterize mRNA-protein interactions in cell extract [15]. Multicolor excitation is needed to monitor the composition and activity of these biological systems, making the advantages of avoiding the multi-band-pass dichroic obvious.

However, the increases in SNR that mmTIRF provides come at the price of significantly increased optical complexity, not only due to the introduction of the micromirrors themselves, but because of the additional optics required to exploit them fully. In objective-type TIRF in general, slight misalignments of the TIRF field inside the high-NA objective can result in significant scatter [16]. mmTIRF provides detailed control over the alignment of the TIRF beam, and the final image quality depends strongly on the details of the optics and optomechanics used to control it. These critical degrees of freedom are fundamentally coupled: adjustments to one affect the others, and all of them affect the position, angle, and uniformity of the excitation beam, which determines the quality of the TIR excitation intensity profile in the sample, or TIRF footprint. The ultimate goal of this manuscript is to show how to precisely, quantitatively control the quality of the TIRF footprint—its position, radial extent, and penetration depth—using the coupled degrees of freedom provided by the mmTIRF system.

mmTIRF systems can be home-built or partially commercial (via the purchase of micromirrors along with a stage designed to support and control them). Helpful protocols have already been published about the construction and alignment of mmTIRF [8, 17, 18] as well as data analysis for CoSMoS [19-21] and single-particle tracking [18]. We refer the reader to them for an overview of the initial alignment of these systems. Our goal in this manuscript is to provide a guide to the optimization of mmTIRF’s sensitive optics to achieve the best imaging performance it can enable. We have used the commercial mmTIRF system from Mad City Labs, which facilitates all the adjustments we will describe throughout the text, especially for the micromirrors. We will review the principles of the design of micromirror-based TIRF, including both the excitation and detection pathways. We will explore the coupling of the critical optical elements and illustrate the effect of small changes in the alignment of four key elements of the system (the TIRF lens, input micromirror, back aperture iris, and combining dichroic). Throughout, we will offer advice on best design, alignment, and maintenance practices.

## 2. Methods

### 2.1 Coupling of the excitation and detection pathways: the central role of the objective lens

In objective-type TIRF, the objective lens is the critical element for both excitation and imaging, so these two pathways are fundamentally coupled. The existing protocols on the alignment of mmTIRF [8, 17] are designed to satisfy three essential criteria relative to the objective lens. First, the excitation lasers must be parallel to the optical axis of the objective, and each must be focussed on its back focal plane (BFP). Second, to attain TIR angles of excitation, it must be possible to radially translate the focal points of these beams across the objective’s BFP. And third, the detection optics must correctly define the imaging plane.

Figure 1a highlights the most important optical components of the excitation pathway. Proceeding along the optical path (counter-clockwise from the bottom of the image), a row of combining dichroics provide independent control of each collimated excitation laser, co-aligning them on the optic axis. A periscope raises the beams to the level of the micromirrors and controls their alignment through the TIRF lens.

The TIRF lens focuses the beams in the back focal plane (BFP) of the objective and onto the input micromirror, which deflects the beams upward into the periphery of the objective but parallel to its axis. The total of six degrees of freedom on the TIRF lens and input micromirror (the XY kinematic knobs on the TIRF lens, and three set screws and axial position of the micromirror) control the exact position and angle at which the excitation beams enter the objective (NA 1.49), which defines the location and quality of the TIRF footprint. The input micromirror is shown from both the side and the “front” (reflective face) to illustrate the positions of the three tip-tilt screws under its base. Two irises are included in the schematic: I_exc_ prior to the TIRF lens controls the size of the TIRF footprint, and I_BA_ beneath the micromirrors controls the effective size of the back aperture and depth of field of the image.

The excitation pathway is aligned to focus the excitation lasers at the edge of the back aperture of the objective while propagating parallel to the optic axis. This alignment is required so the TIRF beam is collimated at the glass-water sample interface and the angle is precisely controllable. Misalignment will cause the footprint to be formed off the objective’s optic axis, which will ultimately make optimal imaging impossible, no matter the perfection of the detection optics.

Incorrect alignment of the detection pathway can also make it impossible to image optimally in the focal plane, even if the excitation optics are perfect. If the detection pathway is misaligned—and in particular if the final imaging lens is not positioned correctly—then a different plane, either slightly above or slightly below the objective’s focal plane, is actually conjugate to the camera. Imaging above or below the objective’s front focal plane induces many of the same effects as misaligning the excitation laser. In particular, it will degrade the SNR and cause the TIRF footprint to form off the optic axis.

In micromirror TIRF, the excitation/detection loop is closed eloquently and visually through the use of the micromirrors. When the interface between the coverslip top surface and sample solution is in the objective’s front focal plane, the TIRF footprint will be centered on the optic axis, and the reflected beam will be centered on the output micromirror (left of Figure 1a). The reflected beam, visible both on the output micromirror and in the back aperture of the objective (or on a white paper held below the objective), is the hallmark of TIRF. When a correctly aligned microscope is in TIRF, the front focal plane is also in focus. Once this initial alignment has been established [17], the input micromirror and the TIRF lens can be adjusted finely for each experiment to compensate for small deviations introduced by the sample and find the best SNR.

In this section, we will discuss each major stage of the mmTIRF microscope—the decoupled optics that control each individual laser, the coupled optics that control the formation of TIRF, and the detection optics—and offer advice for the fine-tuning of the alignment at each stage. We assume the reader is already familiar with the general construction of a mmTIRF microscope, and instead focus on the theoretical and practical concerns that will enable it to be used to its highest potential.

### 2.2 Decoupled (pre-periscope) optics

At least two excitation wavelengths are required for micromirror TIRF to be worth the additional complexity relative to dichroic TIRF. The additional degrees of freedom controlling the micromirrors and optics enable multiple lasers to be optimized in TIRF without loss of signal through a multi-band-pass dichroic. The critical optics for generating the TIRF beam—the micromirrors and the TIRF lens—affect all lasers simultaneously. However, the quality of TIRF depends greatly on the alignment and quality of the excitation beams before they reach this stage of the microscope. Therefore, we advise that every excitation laser should have its own optics for beam expansion, collimation, and filtering before the lasers are coaligned, as at the bottom of Figure 1a.

A schematic of the optical system that expands and shapes each excitation beam is shown in Figure 1b. To generate a large excitation spot that will fill the back aperture of the objective (leading to a large, collimated TIRF footprint in the focal plane, as in Figure 1c), each beam is expanded and collimated with a 10x Keplerian beam expander, combining a 25 mm focal length aspheric lens for focusing the raw laser output and a 250 mm focal length lens for collimation. Adjustable irises to radially “clip” each beam such that only the more uniform “top hat” of each resulting Gaussian intensity profile is used for excitation. This is restricted to a diameter of 6-8 mm by an iris directly in front of the TIRF lens, I_exc_ in Figure 1a, so the final TIRF excitation spot is as close to a top-hat function as possible, providing approximately uniform illumination across the field of view.

To achieve nearly uniform illumination in the center of a beam profile, the initial profile must be a 2D Gaussian. However, many diode lasers can have irregular beam shapes, and most lasers have contributions from higher-order modes, all of which will translate to irregular interference fringes in the TIRF footprint[16]. The beam expander can be used to spatially filter the laser if a pinhole (diameter = 20-50 μm) is placed in the Fourier plane (the shared focal point of the two lenses) to act as a long pass spatial filter.

The other critical element of the beam expander is that the output beam must be precisely collimated. If it is slightly converging or diverging, it will not precisely focus in the back focal plane (BFP), resulting in a TIRF footprint that is not collimated at the FFP and is instead incident at the sample interface at a cone of angles, inducing a high degree of scattering. Great care should be taken to ensure that the excitation spots after the beam expanders are perfectly collimated; a shear interferometer is extremely helpful in achieving this. Alternatively, the beams can be collimated initially “by eye,” and final collimation can be done when each is focused on the BFP of the objective lens. Slight adjustments of the axial position of each collimating lens can achieve as close to identical Airy patterns for each line emerging from the objective lens as possible. This method makes it straightforward to correct any chromatic aberration between the excitation beams, ensuring that each is focused on the BFP as well as possible.

A series of combining long-pass dichroics coalign the lasers precisely along the optic axis prior to the TIRF optics. We advise the use of kinematic mounts with three degrees of freedom (for example, Thorlabs KC1T or KCB1C) to perform very fine alignment at this stage.

Dichroics can introduce aberrations if they are not perfectly flat, or if the mount introduces strain. To avoid this, we recommend using ultraflat, 2-3 mm thick dichroics and—particularly because the thick dichroics introduce additional strain when clamped in conventional mounts—replacing the clamps with an adhesive like epoxy or the milder Picodent. To avoid damage to the optical element, we recommend applying a thin layer of adhesive to the holder, then resting the optic in place and allowing gravity to hold it while the adhesive sets.

### 2.3 Coupled optics: the TIRF lens, input micromirror, and back aperture iris

Figure1c schematically illustrates the paths different rays take through (a simplified view of) the objective barrel when the excitation beam is correctly aligned for TIRF: parallel to the optic axis and focused in the BFP. The wide, focusing beam can be thought of as separate “wedges” (also shown in Figure1c) that emerge from the front focal reference plane (FFRP; the imaginary plane one focal length away from the objective’s front focal plane) collimated and propagating parallel to each other at different locations. In epifluorescence configuration, this beam would be parallel to and centered on the optic axis. In TIRF, they emerge from the FFRP at the critical angle and displaced from the optic axis. The TIRF illumination footprint forms directly on the objective’s optic axis when the sample plane is in focus.

The small “wedge” of illumination highlighted in Figure1c can be used to visualize the consequences of a slight misalignment. Imagining the wedge as an entire, narrower excitation beam illustrates that a slight deviation from vertical in the BFP results in a displacement of the TIRF footprint from the center of the field of view.

A slight defocus in the image (Figure 1d) causes a similar displacement of the TIRF footprint, as the angled illumination follows a slightly longer or shorter path to the interface. If the detection optics are perfectly aligned, ordinary defocus is easy to identify and correct. The danger, as we will see in section 2.4, is that an improperly aligned detection pathway will form an image of a plane other than the true focal plane, and the displacement of the TIRF footprint may be mistaken for excitation pathway misalignment. In that case, a user would incorrectly “realign” the TIRF footprint to the center of the field of view, misaligning it inside the objective and eroding its quality. For this reason, accurate initial alignment of the coupled optics that affect TIRF and accurate positioning of the final image-formation lens are critical to avoid later mistakes in interpreting the TIRF footprint position.

When we refer to the “coupled optics” that affect TIRF, we are primarily referring to three optical elements that control the path of the TIRF beam inside the objective: the TIRF lens, the input micromirror, and the back aperture iris. However, our ability to use these optics for fine adjustments presupposes perfect expansion, collimation, and alignment of the excitation beams, which we discussed in the previous section. The optics that are first encountered by all the excitation lasers play a similar role. For example, two periscope mirrors raise the excitation beams from the height of the table to that of the micromirrors, which are generally about 25 cm up to provide room for the first optics of the detection pathway. Assuming the lasers have been individually well-aligned to the optic axis, the periscope mirrors should be used to maintain this alignment and ensure all the lasers are precisely centered on and normal to the TIRF lens.

The TIRF lens and input micromirror are together responsible for TIRF beam alignment. They ensure correct alignment inside the objective by defining the orientation, angle, and focal position of the TIRF beam. The reviews of mmTIRF construction and alignment already available [8, 17] provide a good guide to the initial alignment of these optics, particularly strategies to ensure that the TIRF lens focuses in the BFP of the objective and the input micromirror is initially aligned to the optic axis of the objective, so we will not retrace these details here. However, every experimental sample introduces small deflections into the imaging system, so the coupled elements should be systematically and iteratively adjusted with every use to restore perfect reflection at the critical angle in a new sample.

The TIRF lens serves multiple important functions. First, in combination with the objective lens, it defines the degree to which the excitation beam is de-magnified at the sample (de-magnification =*f*_obj_ / *f*_TIR_; in our case, 45x de-magnification results from *f*_obj_ = 3.33 mm and *f*_TIR_ = 150 mm). Second, it focuses the excitation beam on the back focal plane (BFP) of the objective lens. Independent control of the focus of each wavelength is achieved with small axial adjustments of the collimating lens in each beam expander to correct for slight chromatic aberrations in the excitation lines, as explained in the section above. Third, as the beam is focusing, its profile becomes small enough to be fully reflected off the top of the entry micromirror, which is positioned at the back aperture of the objective lens to occlude as little of this as possible with still supporting TIR. And finally, the TIRF lens enables two critical adjustments of the beams: it can be adjusted axially to control beam focus, and in X and Y orthogonal to the optical axis for fine beam-angle adjustments.

The final “coupled optic,” positioned just below the micromirrors, is the back aperture iris, which should be sufficiently restricted to block scatter from the micromirrors but not so narrow that it erodes the SNR. Because the iris effectively acts as an aperture stop, it also affects the depth of focus of the field of view, making it one of the critical elements determining the final quality of the TIRF image alongside the TIRF lens and input micromirror.

The output micromirror is placed symmetrically with the input micromirror to catch the totally internally reflected beam and direct it away from the detection optics, and both micromirrors are positioned to leave the maximum possible amount of the back aperture clear for detection. Because the micromirrors occlude part of the back aperture of the objective, the NA available for imaging is slightly smaller than the NA available for excitation. Users should position the micromirrors so the TIRF beam strikes near the top of the reflective face, maximizing the clear space below the objective [18], and bear the slight reduction in detection NA in mind when selecting an objective. The back aperture diameter d_BA_ = 2NA*f*, so for Nikon’s 60x (*f* = 3.33 mm) and 100x (*f* = 2 mm) objectives with NA 1.49, d_BA,60x_ = 9.92 mm and d_BA,100x_ = 5.96 mm. The “slice” of the outer ring of the back aperture that is lost to the micromirrors is a smaller fraction of the 60x objective’s total back aperture diameter. For this reason, we recommend using a 60x objective with a larger back aperture. (It is also worth noting that a 60x objective “de-magnifies” the TIRF footprint less than a 100x objective does, as discussed above, resulting in a larger excitation area.)

### 2.4 Optical elements in the image formation pathway

Although we have deferred discussion of the imaging pathway to the end of this section, arguably the objective, imaging lenses, number of lasers, and the number and layout of the detection channels are the first design elements to establish. Typical CoSMoS microscopes have from two to four lasers (hence detection channels), and the channels are distributed on the halves or quarters of one or two cameras. Here, we give details for an imaging system with four channels arranged in the quadrants of a large sCMOS camera chip (Figure1e); the same principles apply to a detection pathway with two cameras.

The imaging pathway shown in Figure 1e includes three lenses: a tube lens L_1_, which forms the primary image at plane P_1_, and a pair of lenses, L_2_ and L_3_, which form a 4*f* system relaying the image in P_1_ to the final imaging plane, P_2_, on the camera sensor. The final magnification of the system, given at the top of the figure, is defined by the objective and the focal length ratios of the three lenses.

TIRF objectives (apochromat, NA = 1.49) usually have either 100x or 60x magnification. As we mentioned in section 2.3, we recommend using a 60x objective with a large back aperture to maximize the available NA and TIRF footprint size, then adjusting the magnification within the imaging pathway if desired. Objectives are designed to achieve their stated magnification (M_0_ = 60, 100) with a tube lens of a particular focal length (*f* = 200 mm for most Nikon objectives and 180 mm for Olympus), so the initial magnification is M_0_ times the ratio L_1_/L_0_, the tube lens focal length over the design focal length. The 4*f* relay system increases (or decreases) the magnification by an additional factor of L_3_/L_2_. In our system, a 60x Nikon objective is used with three 225 mm focal length lenses, so the magnification is 60 × 225/200 × 225/225 = 67.5x.

A magnification suits our application (paired with our camera, we achieve 96 nm/pixel). However, we would not be able to increase the magnification much more before the displaced color channels began to be vignetted, or clipped by the edges of our 1” lenses. A user who wants to increase the magnification significantly in the detection pathway should consider using 2” optics.

As we discussed in section 2.1, the process of optically conjugating the sample plane to the camera detector is crucial not only for optimal image quality but for accurate formation of the TIRF footprint. To ensure correct conjugation, the detection pathway is best assembled in reverse, from the camera back to the tube lens. The first—and most important individual—step is the accurate positioning of lens L_3_ one focal length away from the camera sensor. As mentioned in section 2.1, this ensures that each channel will be focused at the detector only when light in that channel is collimated prior to L_3_. It is important to take into account how far recessed the sensor is in its casing so the L_3_ lens can be placed to within a mm of accuracy. A typical camera C-mount places the detector 0.69” or 17.5 mm inside the front plate of the camera.

Once L_3_ is placed, the L_2_ lenses can be positioned to conjugate the P_1_ iris plane to the camera detector in all channels. Illumination from a white light (e.g. an LED or phone flashlight) above the objective lens will be split into all the imaging channels so they can be imaged on the detector simultaneously. The channels will not be initially well separated or focused. Each channel’s L_2_ should be adjusted axially to bring the edges of the iris into focus, and the kinematic steering of each channel can be used to position it in its desired location on the detector. Once all the channels are in focus and separated appropriately, the primary image plane (P_1_) is conjugate to the detector in each channel, and the imaging pathway is close to correct alignment.

The tube lens L_1_ should be placed as close as possible to its focal length away from P_1_ so only collimated light emerging from the objective will be focused at P_1_, which is now conjugate with the detector. The only remaining step is then to make the sample plane itself conjugate to P_1_, which is to say, to place it in the front focal plane of the objective. A bright reference sample (e.g. broadband fluorescent microbeads) can be used to check that all the channels are simultaneously in focus; if not, the L_2_ lenses in the out of focus channels can be adjusted axially until this is achieved.

In Figure 1e, the relay system is color-coded to distinguish between regions where the light is converging, diverging, or collimated. The beam is diverging for part of the path, between P_1_ and L_2_. In principle, a single L_2_ could be placed in a long arm prior to the channel separation pathway to avoid any dichroic interfaces with diverging beams, which do contribute to the accumulation of aberrations, as will be discussed below. In practice, we choose to place a separate L_2_ inside each arm to allow maximum independent control over the focus and alignment of each channel and the ability to account for chromatic aberrations. For the same reason, though, if the color channels are to be divided between two cameras, the splitting dichroic should be placed in one of the regions of the relay system where the beam is collimated (i.e. prior to L_3_, meaning that to minimize aberrations, each camera would have its own L_3_).

If the imaging pathway has four color channels, there may be visible astigmatism in the longest-wavelength channel resulting from repeated transmission through thick 45°-angled dichroics that produce refraction-induced astigmatism. The X components of the transmitted light encounter multiple angled interfaces, while the Y components are parallel to it, leading to an astigmatism in the final image, which is slightly amplified when the incident beam is converging or diverging. This astigmatism cannot be eliminated except by splitting the channels over multiple cameras, but there are strategies for minimizing other aberrations associated with the dichroics, as discussed in section 2.2.

## 3. Results

### Optimization of the coupled elements

Ultimately, the purpose of the fine alignments in the preceding section is to produce single-molecule data with high localization accuracy and high SNR. Although the theoretical goal of aligning the excitation optics is to ensure the excitation beam is parallel to the optic axis, the practical goal is to produce the highest-quality image, and therefore, the highest-quality experimental data. Because every sample introduces different optical conditions—refractive index, coverslip geometry, and even surface preparation—the final alignment changes with every sample, and the coupled elements have to be adjusted to find the optimal alignment. These small changes to the input micromirror, TIRF lens, and iris have a dramatic impact on image quality. The goal of this section is to illustrate these effects as a guide to visual alignment.

### 3.1 Input micromirror

Figure 2 shows the effect of small rotations of the micromirror set screws on the TIRF footprint. Starting from optimal alignment (Figure 2a), we adjusted each of the three set screws by 1/8 or 1/4 of a rotation clockwise and counterclockwise. A schematic of the set screw locations under the input micromirror is shown in Figure 2b. All three set screws affect the position and angle of the micromirror both transverse and parallel to the optic axis, and in each case, adjustments affect both the footprint position and signal quality (SNR/TIRF penetration depth).

**Figure 2.**
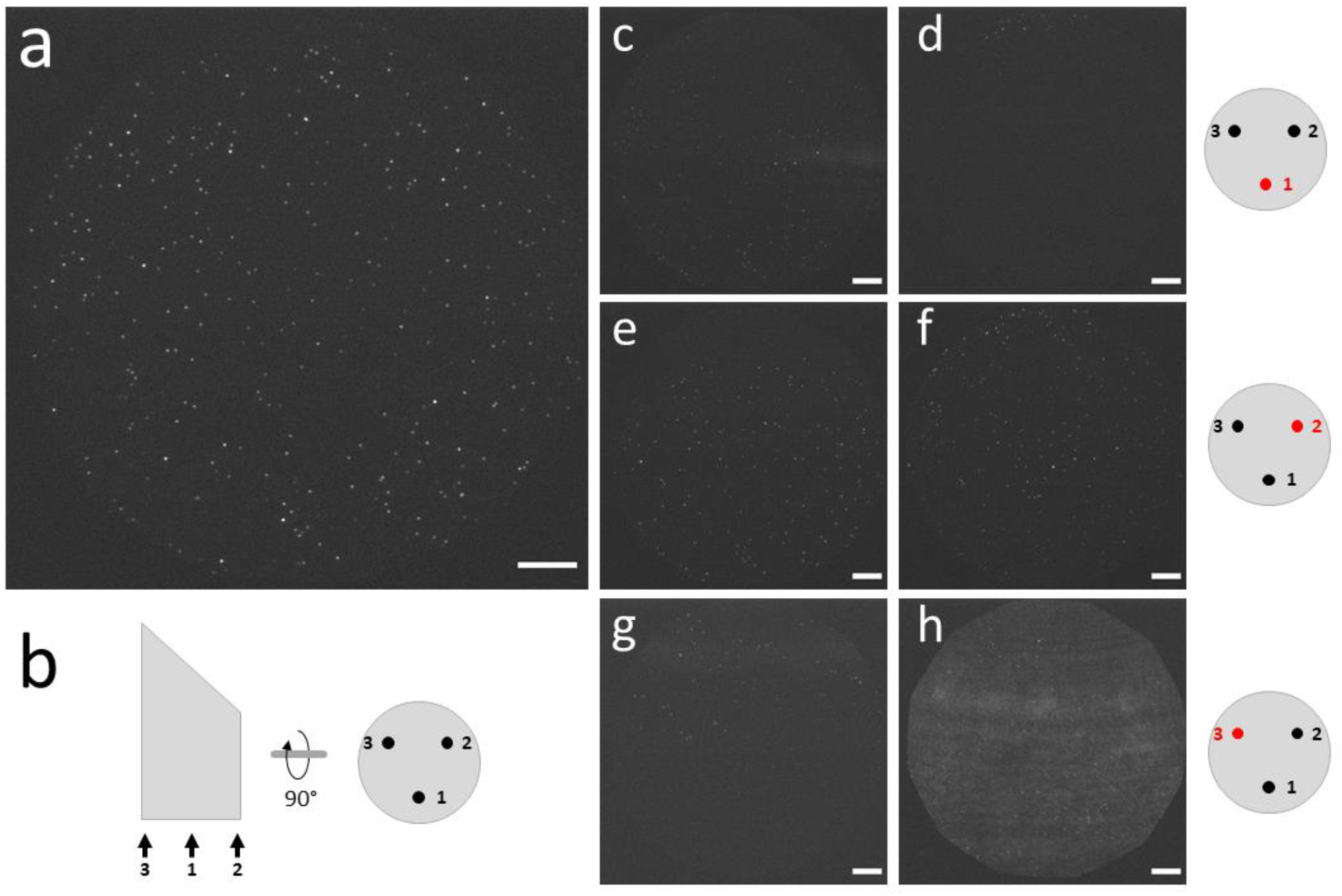
Effect of input micromirror alignment on image quality. (a) Cy5-labeled streptavidin illuminated by a 638 nm laser at 12 mW in optimal TIRF conditions. (b) Schematic of the input micromirror with the locations of set screws 1-3 labeled. (c-h) The same field of view (FoV) after rotating the following set screws: set screw 1, 1/8 of a turn counterclockwise (c) and clockwise (d); set screw 2, 1/4 of a turn counterclockwise (e) and clockwise (f); set screw 3, 1/8 of a turn counterclockwise (g) and clockwise (h). Schematics to the right highlight the relevant set screw in red. All scale bars are 10 μm.

The effect of rotating set screw 1 by 1/8 of a turn is shown in Figure 2c-d. A counterclockwise rotation (Figure 2c) tips the footprint primarily down in the camera field of view (FoV) (which in our case is parallel to the optic axis), as well as degrading the SNR (the mean signal divided by the standard deviation of the background) from 17.8 to 8.5 and introducing some scatter (refraction bands at the right of Figure 2c). A clockwise rotation of the same magnitude displaces the TIRF footprint almost entirely out of the FoV (only a small number of dim PSFs are visible at the top of Figure 2d). A rotation of 1/8 turn corresponds to tightening/loosening the screw by 0.1 mm, highlighting the extreme sensitivity of the micromirrors and the dramatic effect small adjustments can have.

Figure 2e-h show the effects of counterclockwise and clockwise rotations of set screws 2 and 3 on the TIRF footprint. A slightly larger rotation of set screw 2 (1/4 rotation) deflects the TIRF footprint diagonally in the field of view, while 1/8 rotations of set screw 3 primarily change the image quality by introducing a large tilt parallel to the optic axis, directly walking the component of the angle of incidence parallel to the optic axis away from the critical angle. In Figure 2g, a counterclockwise deflection decreased the SNR compared to Figure 2a by a factor of 1.8, while in Figure 2h, a clockwise deflection decreased the SNR by a factor of 7.7, primarily because of the obvious increase in scatter.

### 3.2 TIRF lens

The kinematic degrees of freedom (DoFs) that control the lateral positioning of the TIRF lens are easier to intuitively map onto deflections of the TIRF footprint. In Figure 3a-b, 1.6 mm translations in the X direction (two turns of the kinematic knob) deflect the beam left or right of the optic axis. For small enough deflections, the component of the incident angle parallel to the optic axis is approximately the same, so the primary effect is horizontal deflection of the footprint, although the SNR also visibly decreases between full alignment (shown in Figure 2a for the same FoV) and clockwise rotation (Figure 3b).

**Figure 3.**
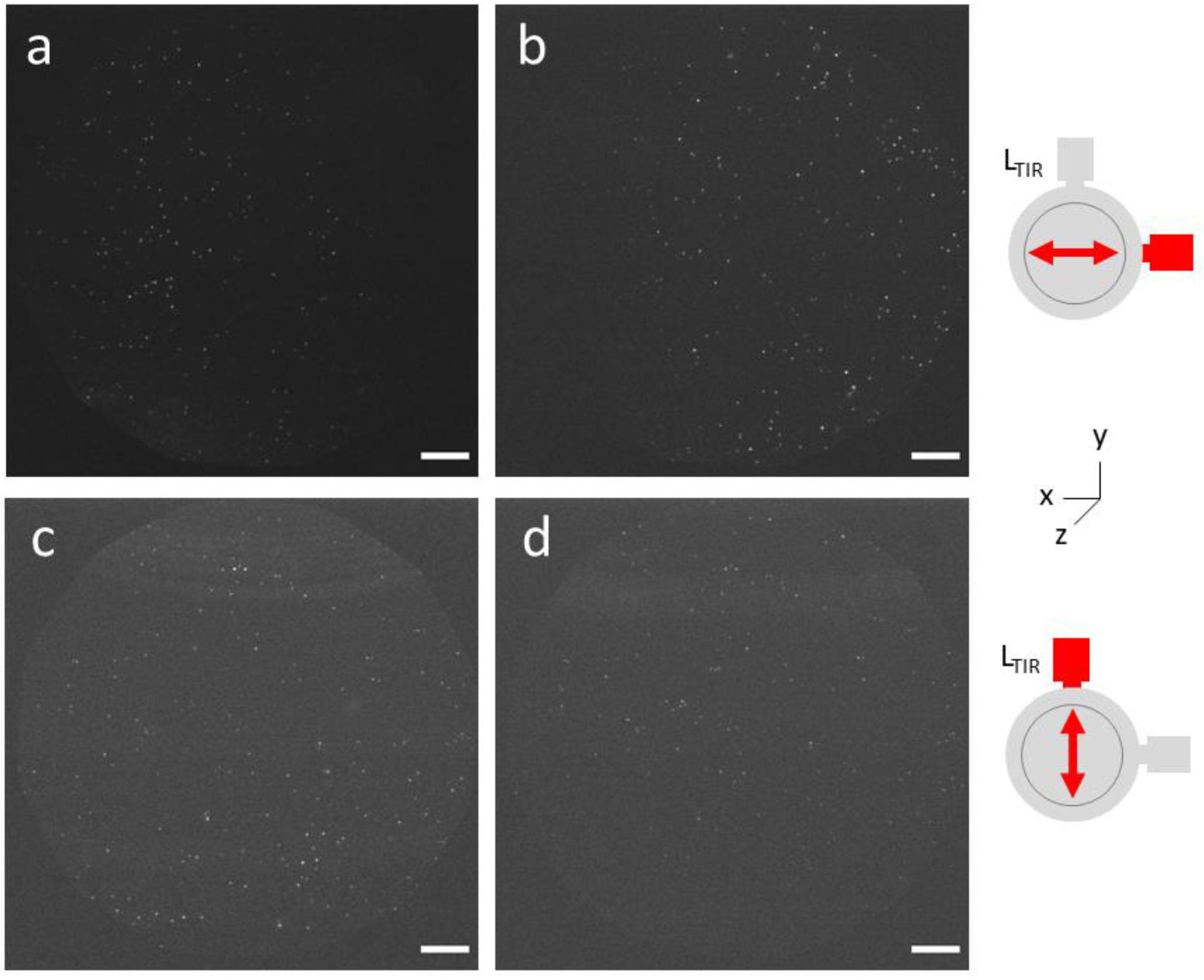
Effect of TIRF lens (L_TIR_) alignment on image quality. The same FoV is shown under optimized imaging conditions in Figure 2a. (a-b) The FoV after turning the X kinematic screw 2 turns counterclockwise (a) or clockwise (b), for a total translation of ±1.6 mm. (c-d) The same FoV after turning the Y kinematic screw 1 turn counterclockwise (c) or clockwise (d), for a total translation of ±0.8 mm. The schematics highlight the relevant kinematic positioner and direction of movement in red. The axes illustrate how L_TIR_ as shown here aligns with its representation in Figure 1a. All scale bars are 10 μm.

Translations in the Y direction (Figure 3c-d) deflect the beam above or below the optic axis prior to the objective (the z direction in Figure 1a), which translates directly into a deviation from the TIRF angle. In Figure 3c-d, the excitation beam does appear to move slightly up and down in the FoV following a 0.8 mm translation, but the primary effect is clearly the degradation of the SNR away from the critical angle (by a factor of 2.3 in the counterclockwise direction and 2.8 in the clockwise direction).

### 3.3 Back aperture iris

The back aperture iris is not present in dichroic TIRF. Micromirror TIRF requires it to exclude the micromirrors from the effective back aperture of the objective, and it has a subtle but profound effect on the overall imaging quality of the system. Figure 4(a, b, c)-i show a field of view containing bright, 0.1 μm-diameter Tetraspeck calibration beads with the micromirrors and TIRF lens in their optimal positions, while the iris is either over-restricted by 1/8 of a turn (about 3 mm change in diameter; Figure 4a), optimized (about 7.5 mm diameter on our setup; Figure 4b), or opened by 1/8 of a turn (Figure 4c). The effect of an over-restricted back aperture on the SNR (Figure 4a-i) is obvious: too much fluorescence is excluded, and the signal is greatly reduced. However, opening the iris too much also causes image degradation: although the SNR in Figure 4b-i (optimal) and Figure 4c-i (open) initially appears similar, there is elevated background noise in Figure 4c-i, as scatter is now permitted into the imaging pathway.

**Figure 4.**
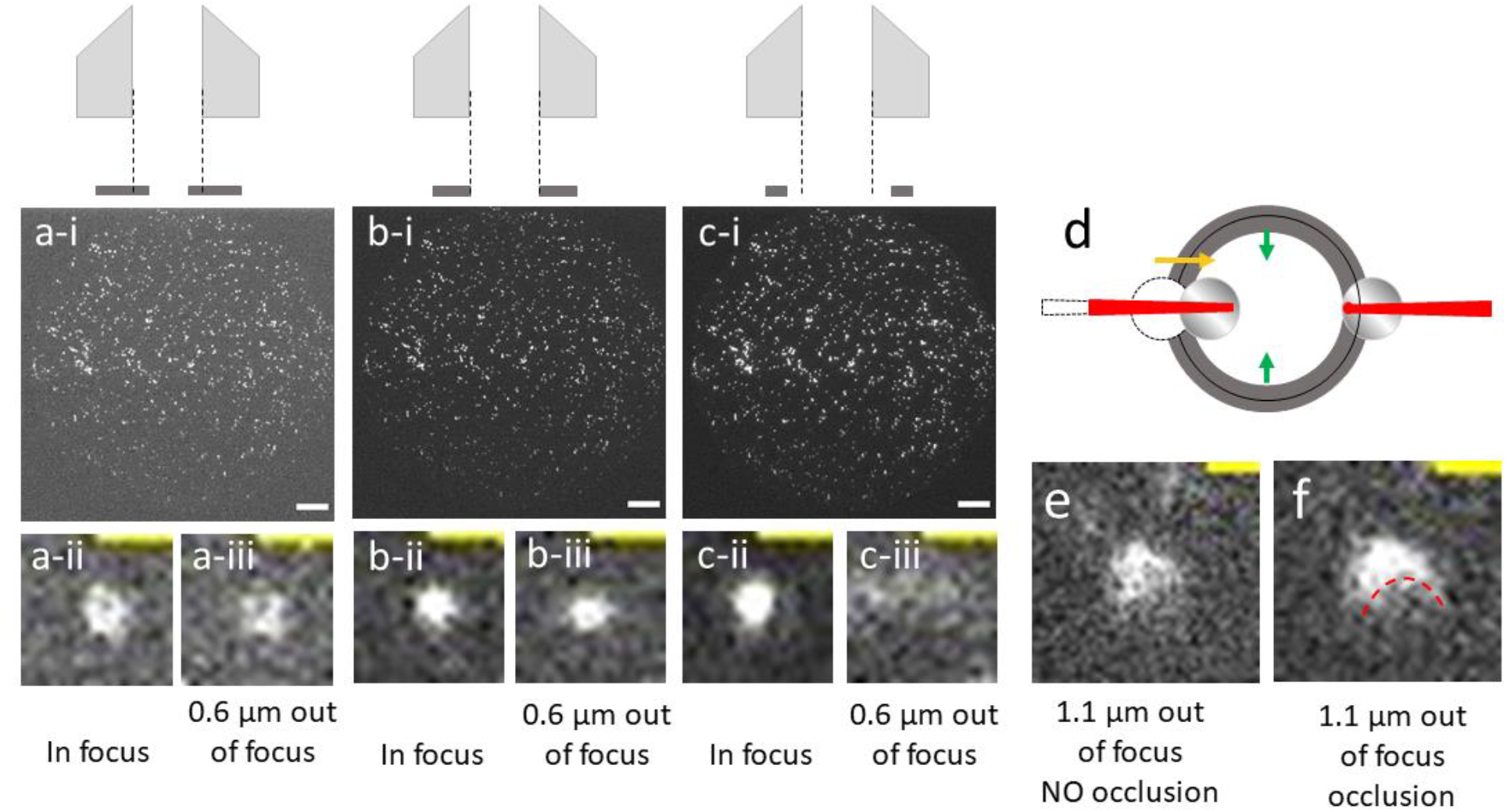
Effect of iris diameter and micromirror position on image quality. 0.1 μm Tetraspeck beads were imaged with a 638 nm laser at 1 mW. (a-i) The effect of closing the back aperture iris 1/8 of a rotation past optimal (schematically illustrated above). (a-ii) Close view of a PSF in focus with the iris over-restricted, and (a-iii) the same PSF 0.6 μm out of focus. (b-i) Image of the FoV with optimal iris diameter. (b-ii) Close view of the same PSF in focus and (b-iii) 0.6 μm out of focus. (c-i) The effect of opening the iris 1/8 of a rotation beyond optimal. (c-ii) Close view of the same PSF in focus and (c-iii) 0.6 μm out of focus. (d) Schematic of the micromirrors and iris viewed from above, illustrating a translation of the output micromirror (left) into the back aperture. (e) Close view of a PSF illuminated at 488 nm and 3.5 mW, 1.1 μm out of focus with the micromirror in its optimal position. (f) Close view of the same PSF, 1.1 μm defocused with the micromirror partially occluding the aperture. The red dashed line emphasizes the location of the micromirror’s shadow, distorting the profile of the PSF. White scale bars in panels -i are 10 μm. Yellow scale bars in panels -ii and -iii are 1 μm.

One might infer from this that the correct way to optimize the iris diameter is to find the best SNR that does not introduce scatter. However, the iris has a second and more subtle impact on PSF quality, which should also be considered. Figure 4(a, b, c)-ii and -iii highlight a particular PSF in the center of the FoV and illustrate the effect of the iris diameter on PSF shape and quality, both in focus and 0.6 μm away from the focal plane. When the iris diameter is over-restricted (Figure 4a-ii, iii), the SNR is reduced as before, but the PSF retains a symmetrical shape and comparable width when the image is slightly out of focus (in Figure 4a-ii, σ_x_ = 276 nm, σ_y_ = 305 nm; in Figure 4a-iii, σ_x_ = 296 nm, σ_y_ = 292 nm). In contrast, when the iris is open (Figure 4c-ii, iii), both scatter and out-of-focus rays collected far from the objective’s optic axis are collected, so the PSF diverges very rapidly away from the focal plane. An overly-open iris effectively decreases the system’s depth of focus and degrades the quality of the PSFs detected.

The goal when optimizing the diameter of the back aperture iris is shown in the center of Figure 4b. The ideal diameter actually strikes the best middle ground between a high SNR and acceptable depth of field and PSF quality: the defocused PSF in Figure 4b-iii has a similar width (σ_x_ = 239 nm, σ_y_ = 292 nm) to the PSF in Figure 4a-iii, but with much improved SNR, 29.6 across the optimal field of view in Figure 4b compared with 15.4 with the iris closed in Figure 4a.

### 3.4 Output micromirror positioning

The position of the output micromirror also affects PSF quality. The output micromirror should be symmetrically positioned with the input micromirror relative to the objective. Visual inspection of the two micromirrors from the side is an excellent way to quickly assess the quality of their alignment: any deviation from symmetry in height, angle, or displacement from the optic axis is a sure indicator of misalignment of one or both mirrors.

The consequences of one specific output micromirror misaligment are shown in Figure 4d-f. If the output micromirror is too close to the optic axis, as shown in the schematic (Figure 4d), it will occlude the back aperture and interfere with the collection of the fluorescence signal. Figure 4e shows a point spread function 1.1 μm away from the focal plane in the absence of occlusion. Figure 4f shows the same PSF with occlusion by the micromirror, visible in the form a distinct “shadow” obscuring part of the PSF’s profile.

The effects on PSF quality in sections 3.3 and 3.4 are subtle and hard to visually detect in a focused image with bright, apparently homogeneous PSFs, so it is tempting to discount their importance. However, PSFs of high quality and uniformity with high photon counts are essential to most applications of CoSMoS, which depend on highly accurate localization of individual fluorophores. If the depth of field is shallow due to a misaligned iris (section 3.3 and Figure 4a-i), slight defocus will severely degrade the PSF width and photon count even if the TIRF footprint has been optimized, decreasing the localization accuracy of the imaging system as a whole. Likewise, deviations of the PSF from the 2D normal distribution decrease the accuracy of localization fitting, even when SNR and photon counts are high. The iris and output micromirror must be correctly positioned in order to take full advantage of the SNR achieved by optimizing the TIRF beam angle.

### 3.5 Effect of a deliberate misalignment

To demonstrate the sensitivity and coupling of the optics discussed in the preceding sections, we deliberately misaligned one of our excitation lasers using its combining dichroic, then attempted to compensate using the input micromirror and TIRF lens (Figure 5). Far from recovering the image quality, this has the effect of propagating misalignment throughout the system, with the result that none of the lasers were able to form a well-aligned TIRF footprint, even the two that were initially well aligned.

**Figure 5.**
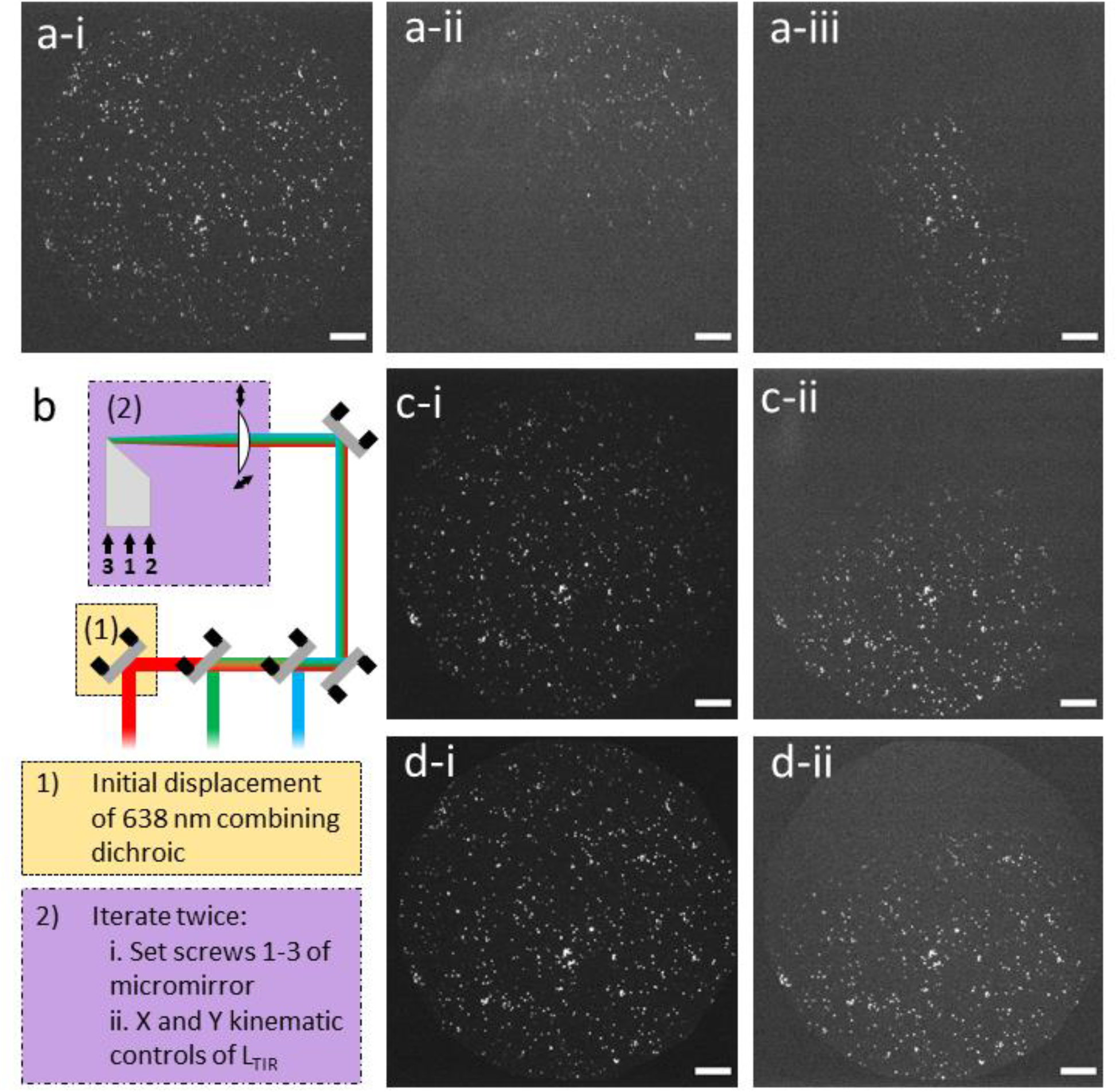
Effect of a misaligned input optic on global TIRF alignment. 0.1 μm Tetraspeck beads were separately imaged at 488 nm, 532 nm, and 638 nm (3, 0.7, and 0.1 mW, respectively). (a-i) Initial state of the FoV under red excitation before misalignment. (a-ii) Result of initial displacement of the red laser. (a-iii) Result of attempting to realign the red laser using the coupled optics. (b) Schematic of the misalignment and realignment process. Box (1) (yellow): initial displacement of the 638 nm combining dichroic. Box (2) (purple): summary of steps taken to realign the red laser using the input micromirror and TIRF lens. (c-i) Initial state of the FoV under green excitation. (c-ii) Final alignment state of the green laser after realigning based on the red laser. (d-i) Initial state of the FoV under blue excitation. (d-ii) Final alignment of the blue laser after realigning based on the red laser. All scale bars are 10 μm.

Figure 5a-i shows a field of 0.1 μm Tetraspeck beads illuminated at 638 nm and imaged in the red detection channel of our camera. As shown in the schematic in Figure 5b, we chose to misalign the red laser using the DoFs of its combining dichroic, with the result shown in Figure 5a-ii. Our excitation lasers are combined in order of decreasing wavelength using long-pass dichroics, so we chose to misalign red to avoid any effect of the slight change in interface angle on the other lasers.

We then attempted to “realign” the red laser using the coupled optics discussed above (the input micromirror and TIRF lens; we did not change the back aperture iris diameter). Our best result, shown in panel a-iii, was achieved by iterating twice through five degrees of freedom—the three micromirror tip/tilt screws and the two TIRF lens kinematic positioners—as highlighted in purple in panel b. None of the DoFs were moved by more than 1/4 of a turn. Further adjustment beyond two iterations only degraded the image quality further, so we worked backwards to restore the final image shown in Figure 5a-iii.

The red laser’s TIRF footprint is much smaller and no longer parallel with the optic axis (Y in our field of view). The SNR within the footprint is reduced by a factor of 1.3. We considered that the initial deflection of the red laser caused it to clip on the input iris, but opening it from 7 mm to 10.5 mm slightly elongated the footprint without restoring its lateral distribution.

This demonstration shows that it is impossible to use the fine, coupled DoFs to create a good TIRF beam if the excitation lasers are not initially well aligned with the optic axis. Attempting to do so also displaces and degrades the quality of excitation footprints that were well aligned. in Figure 5c, the SNR in the green channel (illuminated at 532 nm) reduces by a factor of 1.3 after “realignment”; in Figure 5d, the SNR in the blue channel (illuminated at 488 nm) reduces by a factor of 2.0. If a user notices that the TIRF footprint in one channel is worse than that in the other(s), the user should first correct any upstream misalignments. Attempting to optimize using only the coupled DoFs will result in a global misalignment.

## 4. Discussion

### 4.1 Optimal use and maintenance of mmTIRF

We have shown that the critical optics that control TIRF footprint quality in a micromirror system are coupled, and that obtaining high-quality data from the microscope requires an understanding of how they interact. Users who are not fully aware of the intricacies of the coupled optics can slowly degrade the SNR of the system by “walking it away” from optimal alignment through successive small adjustments. In the authors’ experience, overreliance on one optic, like the TIRF lens, while ignoring its coupling to the others can limit the user to suboptimal TIRF conditions and also eventually result in a misalignment that requires global adjustments to fix. At minimum, new users should be trained to perform these daily adjustments. We have found that it is beneficial for a new mmTIRF user to fully align the whole coupled section of the microscope (from the periscope mirrors through the back aperture iris). In this way the user develops an intuition for the effect of each DoF on the image, the small range of movement over which adjustments are necessary, and how to systematically optimize the three coupled DoFs without walking any of them out of alignment.

Users with very different samples and buffers should be aware of the large impact these differences have on the TIRF footprint. Different buffer or coverslip refractive indices will alter the critical angle, and the thickness and roughness of the sample holder will affect the amount of scatter. Users should expect to adjust all the coupled DoFs before each measurement to optimize the SNR in their specific sample. As a corollary to this observation, users are advised to obtain a calibration sample as closely optically matched to their experimental sample as possible (e.g. it should be prepared on coverslips of the same material and thickness, using the same or a very similar buffer). Even with a perfect match, there will still be slight differences between the calibration sample and the real one; minimizing these differences will make the alignment as close as possible.

Although it is rarely discussed in the TIRF literature, we note that surface passivation affects the critical angle as well. The real refractive index at a chemically treated (e.g. PEGylated or otherwise prepared) interface is not precisely n_buffer_/n_glass_. All other conditions being equal, users should expect to optimize their optics even if the surface is the only difference between a pair of samples.

Aside from daily adjustments of the TIRF DoFs, users can reasonably expect to realign the periscope and combining dichroics once every few months. It will become apparent that this is necessary when the excitation beams drift visibly off-center on the excitation iris. In general, any freestanding optic in the system will drift on this timescale, while those secured in multiple dimensions (e.g. using a cage system) will only need to be adjusted annually or less.

When assessing which optics should be realigned, users should take care to distinguish between separate scenarios. If only one laser has drifted, or if multiple lasers are decentered in different directions, the cause is most likely the relaxation of the combining dichroic(s). In these scenarios, only the combining dichroic(s) associated with those lasers should be realigned. However, if all the lasers are decentered in the same direction, they are subject to a coupled misalignment, and the periscope mirrors should be realigned.

When adjusting the combining dichroics, the user should begin at the “front” of the line of dichroics (the shortest wavelength if long-pass mirrors are used) and work systematically back to the last. This ensures that any small deflections that occur when the longest-wavelength lasers pass through the pile of dichroics are fully accounted for during alignment.

### 4.2 Relative advantages and disadvantages of commercial and home-built CoSMoS

As mentioned in the section 1, mmTIRF systems are commercially available. Mad City Labs, Inc., sells the micromirrors along with a stage and support arms for aligning them. The tip-tilt micromirror controls we have described in this paper are the commercial ones. The most obvious advantages of a commercial mmTIRF system are simplicity—it integrates the micromirror support system directly with software to control the nanopositioning stage and optional TIRF lock arm—and precision, as the controls have been pre-calibrated and machined specifically for the purpose.

It is possible, though nontrivial, to assemble an analogous system from individual parts. 45° rod mirrors are commercially available from multiple sources (Edmund Optics and Knight Optical, to name two). The user will want mirrors of length 4-5 mm and diameter 2-3 mm. The most substantial issue the builder of a home-made mmTIRF will encounter is the design of a custom holder for the micromirrors. One of the existing protocols for mmTIRF [17] used a prior version of the commercial system from Mad City Labs, but it describes the details of its construction and alignment and provides a guide to how these holders were constructed (using the Newport MT-XYZ linear stage for control). Another recent publication [18] deals in detail with the construction and alignment of a home-built mmTIRF system for single particle tracking experiments.

We expect that a custom mmTIRF may be beneficial if the user intends to integrate additional specialized optics under the objective; work at wavelengths that require unusual reflective coatings on the micromirrors; use a specialized stage or holder; or thermally isolate the microscope in a way that makes direct access to the optics during experiments impossible. Thermal isolation, for example, may make it desirable to mount the micromirrors such that their tip/tilt controls can be accessed via knobs outside an isolation box.

### 4.3 Conclusions

As colocalization experiments become more complex and incorporate more excitation wavelengths, the dichroic that enables objective-type TIRF becomes a limiting factor. Micromirror TIRF not only avoids the decrease in sensitivity that results from absorption in the multi-band-pass dichroic but provides an unprecedented degree of control over the TIRF footprint. This control gives the experimenter the ability to optimize the image quality—and hence data quality—extremely sensitively.

Making the best use of mmTIRF requires a thorough understanding of the dual role played by the objective in both the excitation and detection pathways. The conditions required for TIRF, which involve both the collimation and alignment of the excitation lasers (section 2.2) and accurate positioning of the imaging optics (section 2.4), are all defined in reference to the objective. The coupling between the elements that control the TIRF beam’s orientation (TIRF lens, input micromirror, back aperture iris) must also be understood to bring the system into a globally optimal alignment (section 2.3), which also depends on the specific sample being used. We find that a thorough grasp of this system is pedagogically useful: a microscopist who takes the time to understand the interactions between its elements will have mastered all the most important concepts relevant to single-molecule imaging. Our goal has been to call attention to the most critical points in this complex and coupled microscopy technique and offer advice on how to ensure that a potentially powerful approach to TIRF is in fact being used to its utmost capacity.

## Acknowledgments

We thank many people who contributed to the design and development of our micromirror CoSMoS instrument: Dr. Eric Drier, Dr. Edo van Veen, Dr. Serge Vincent, Dr. Humberto Sánchez, Dr. Fatemeh Moayed, Dr. Belén Solano, and Dr. Sam Leachman. We particularly thank Dr. Drier for many fruitful discussions about the optimization of micromirror CoSMoS and for his feedback on this manuscript. We also thank Prof. Carlas Smith and Mr. Jelmer Cnossen for pointing out the importance of the PSF profile for localization accuracy. ND acknowledges funding from the Netherlands Organization for Scientific Research (NWO) through the Top grant [grant number 714.017.002], from the Dutch Foundation on Fundamental Research on Matter (part of NWO) [grant number 16PR1047], and from the European Research Council through an Advanced Grant [REPLICHROMA; grant number 789267].

## Author Contributions

**Kaley McCluskey**: Conceptualization; Formal analysis; Resources; Investigation; Methodology; Visualization; Writing – original draft; and Writing – review & editing. **Eric A. Drier**: Conceptualization; Methodology; Resources; and Writing – review & editing. **Nynke H. Dekker**: Conceptualization; Funding acquisition; Resources; Writing – review & editing.

